# Spatiotemporal biases in localization and interception: common underlying mechanisms?

**DOI:** 10.1101/2025.04.25.650604

**Authors:** Anna Schroeger, Simon Merz

**Affiliations:** Department of Experimental Psychology, Justus Liebig University Giessen, Germany; Center for Mind, Brain and Behavior, University of Marburg, Justus Liebig University Giessen and University Darmstadt, Germany; Department of General Psychology, Trier University, Germany; Institute for Cognitive & Affective Neuroscience, Trier University, Germany

## Abstract

Human perception of space, time and motion is subject to several biases. Lab studies showed such effects in psychophysical judgements of location, but also in action-tasks, like predicting motion and intercepting. Given that similar underlying processes have been proposed for some of these biases, we tested for a shared mechanism by correlating them across observers. Using the classical implied motion sequence, participants either indicated the remembered location of an intermittently presented dot consistently ‘moving’ from one location to the next, or intercepted a predicted future location of the same intermittently presented dot. We examined whether the errors in those tasks are associated by correlating i) the overall amount of overshooting, and ii) the effect of temporal manipulations of the ‘jump’ duration on these biases across participants. We found two medium correlations indicating that these two biases are indeed related to each other. Participants who show a larger effect in one task also show a larger effect in the other task, and participants with larger proneness to temporal features show them consistently across both tasks. This suggests a shared underlying mechanism, and theoretical implications are discussed.

## Introduction

In our daily lives, we frequently interact with moving objects, for instance, when catching or hitting a ball or deciding whether to cross the street before a car arrives. We seem to be quite successful in those tasks, even though lab studies revealed a series of temporal and spatial biases in the context of motion perception (Freyd & Finke, 1984; Helson, 1930; Helson & King, 1931). While some areas of research have tried to investigate these biases using simple psychophysical judgment or localization tasks (Fröhlich, 1923; Hubbard, 2005, 2018; Müsseler & Kerzel, 2018), others have focused on the prediction of space, time or motion using action tasks like interception (Brenner & Smeets, 2011; de la Malla et al., 2019; Schroeger et al., 2021).

Interception refers to an action to stop or interfere with a moving object, e.g. hit or catch a ball. Both directions reveal interesting biases, but it is still unclear whether these effects originating from different paradigms are actually caused by the same underlying factor. This study aims to integrate two key approaches in motion research by comparing spatial biases in two similar tasks: one involving spatial localization of moving objects (based on representational momentum) and the other involving spatial prediction of moving objects (leveraging the tau effect).

Regarding the localization of dynamic stimuli, several biases in as well as against the direction of motion have been reported over the years: for example, the fröhlich effect, onset-repulsion effect, offset-repulsion effect, or representational momentum effect (for reviews and discussion, Hubbard, 2018; Merz et al., 2022; Müsseler & Kerzel, 2018). Localization has been investigated across diverse set-ups, such as different sensory modalities (e.g., audition: Getzmann et al., 2004; vision: Hubbard & Bharucha, 1988; touch: Merz, Deller, et al., 2019; Merz, Meyerhoff, et al., 2019); different trajectories (circular: Freyd & Finke, 1984; Merz et al., 2022; linear: Hubbard & Bharucha, 1988) and different stimulus motion types (intermittent presentations of stimuli with changing positions along linear trajectories, i.e. implied motion: Freyd & Finke, 1984; Merz, Meyerhoff, et al., 2019; consistent motion: Hubbard, 1990; Merz, 2022). The final location of a moving object is most commonly misjudged as being shifted in motion direction - a phenomenon called representational momentum (Freyd & Finke, 1984; for reviews, see Hubbard, 2005; Merz et al., 2022). Theoretical accounts trying to explain this effect range from low-level perceptual explanations to memory based effects (for discussion, see Hubbard, 2010).

Similar to stimulus localization, prediction has been investigated across different sensory modalities (vision: for review, see Battaglini & Ghiani, 2021; Fooken et al., 2021; audition: e.g., Schroeger, Raab, et al., 2022), stimulus trajectories (linear: e.g., Alderson & Whiting, 1974; for review, see Tresilian, 1995; simulated ball trajectories: Fooken et al., 2016; Kreyenmeier et al., 2017, 2023; ballistic trjectories including gravity: Delle Monache et al., 2019) and stimulus motion types (intermittent presentations of stimuli with changing positions along linear trajectories, i.e. implied motion: e.g., Schroeger, Grießbach, et al., 2022; Schroeger, Raab, et al., 2022; consistent motion: e.g., Delle Monache et al., 2019; Fooken et al., 2016). Recently, Schroeger, Raab, et al. (2022) observed the so-called tau effect (Benussi, 1913; Helson, 1930; Helson & King, 1931) within such a prediction task that largely resembles a representational momentum paradigm. The tau effect describes the observation that the perceived distance between two objects depends on the temporal delay between their presentations: the longer the delay, the further apart two stimuli are perceived. In line with this, in the prediction-version of this task, Schroeger, Raab et al. (2022) found that participants systematically overshot the predicted location of an intermittently moving target when the temporal delay between presentations was long (but see, Merz et al., 2022, for possible directions of the effect).

Localization and interception tasks for intermittently presented motion share similar paradigms, reveal biases in the same direction and are influenced by temporal features, differing mainly in instructions to remember the final location or predict a future one. This suggests a connection between the tasks, though direct comparisons are scarce in the literature. Previous research has associated representational momentum with time-to-contact estimations (Gray & Thornton, 2001), showing a transfer of localization effects to temporal prediction. To extend this understanding by including direct action, our study aims to analyze both spatial localization and spatiotemporal prediction through two comparable tasks based on representational momentum and the prediction-version of the tau effect.

In the current study, we tested whether biases in localization and interception are indeed interrelated. We applied two highly comparable paradigms of a ‘moving’ dot intermittently presented at 5 or 4 locations across a touchscreen. Participants either indicated the final location by tapping the screen after the presentation ended or they predicted the next location by intercepting at the predicted time in the predicted location. We first tested whether the overall amount of overshooting is related between both tasks by leveraging individual differences in the size of these biases and correlating them across participants. Second, we tested whether both localization and interception performance are affected by manipulations of the temporal intervals between target presentations (‘jumps’). Again, individual differences in the effect sizes of temporal manipulations served to relate localization and interception to each other.

## Methods

### Participants

A power analysis on the correlation between both tasks revealed a number of 67 participants needed to find at least a medium effect (ρ_H1_ = 0.3, α = 0.05, power = 0.8, ρ_H0_ = 0). A total of 67 participants (mean age, male: female:) took part in the experiment. One additional participant was excluded because they confused the task instructions. Participants provided informed consent and received compensation of 8€/h or course credits. The study is part of a project with ethics approval from the local ethic committee (approval number: LEK-2021-0028).

### Design

The experiment was conducted with a 2 (task: localization vs. interception) x 3 (temporal interstimulus intervals: long: 450 ms; medium: 250 ms; short: 50 ms) x 3 (spatial interstimulus intervals: short: 30 px = 1.11 cm; medium: 90 px = 3.32 cm; large: 150 px = 5.53 cm) design.

Please note that the manipulation of spatial interstimulus intervals only served the purpose of introducing variability in the motion trajectories. For each trial, spatial (horizontal as well as vertical location in pixels) as well as temporal (moment of touch) response information were stored.

### Materials

All stimuli were presented on a 32” Display ++ LCD Monitor (1920×1080 px) from Cambridge Research Systems including IR Touchscreen technology. Visual stimuli were black dots (15 px = 0.55 cm) on a grey background. Instructions were standardized and given via the experimental software, PsychoPy version 2023.1.3 (Peirce et al., 2019). Within a trial the stimulus was presented consecutively on 4 (interception task) or 5 (localization task) locations ordered from left to right or right to left, simulating intermittent motion. Participants’ task was to indicate the fifth location of the stimulus. In the localization task, the fifth location was presented, whereas in the interception task, the location of the fifth stimulus had to be temporally and spatially predicted by the participants.

### Procedure

To start a trial, participants were instructed to depress the spacebar and keep it pressed until they wanted to initiate their response in the interception paradigm or until visual presentation was over in the localization task. Following the depress, a fixation cross appeared at the center of the screen for a random interval ranging between 400-600 ms drawn from a uniform distribution. As soon as it disappeared, the target stimulus was shown the first time and then intermittently presented while ‘moving’ horizontally either from right to left or left to right. The target appeared either four (interception task) or five times (localization task), with fixed temporal (long: 450 ms; medium: 250 ms; short: 50 ms) and spatial interstimulus intervals (short: 30 px = 1.11 cm; medium: 90 px = 3.32 cm; large: 150 px = 5.53 cm) and a fixed stimulus duration of 250 ms. Within one trial, the temporal and spatial interstimulus intervals (ISI) were constant but randomly altered between trials. For the interception task, participants needed to spatially and temporally predict the occurrence of the target at the fifth location. For the localization task, participants were allowed to start moving their hand following the disappearance of the target at the fifth location. Each participant conducted both tasks in a blocked manner and used their index finger on the touchscreen (**Fig. 1**). The order of the tasks was counterbalanced across participants.

**Figure 1.**
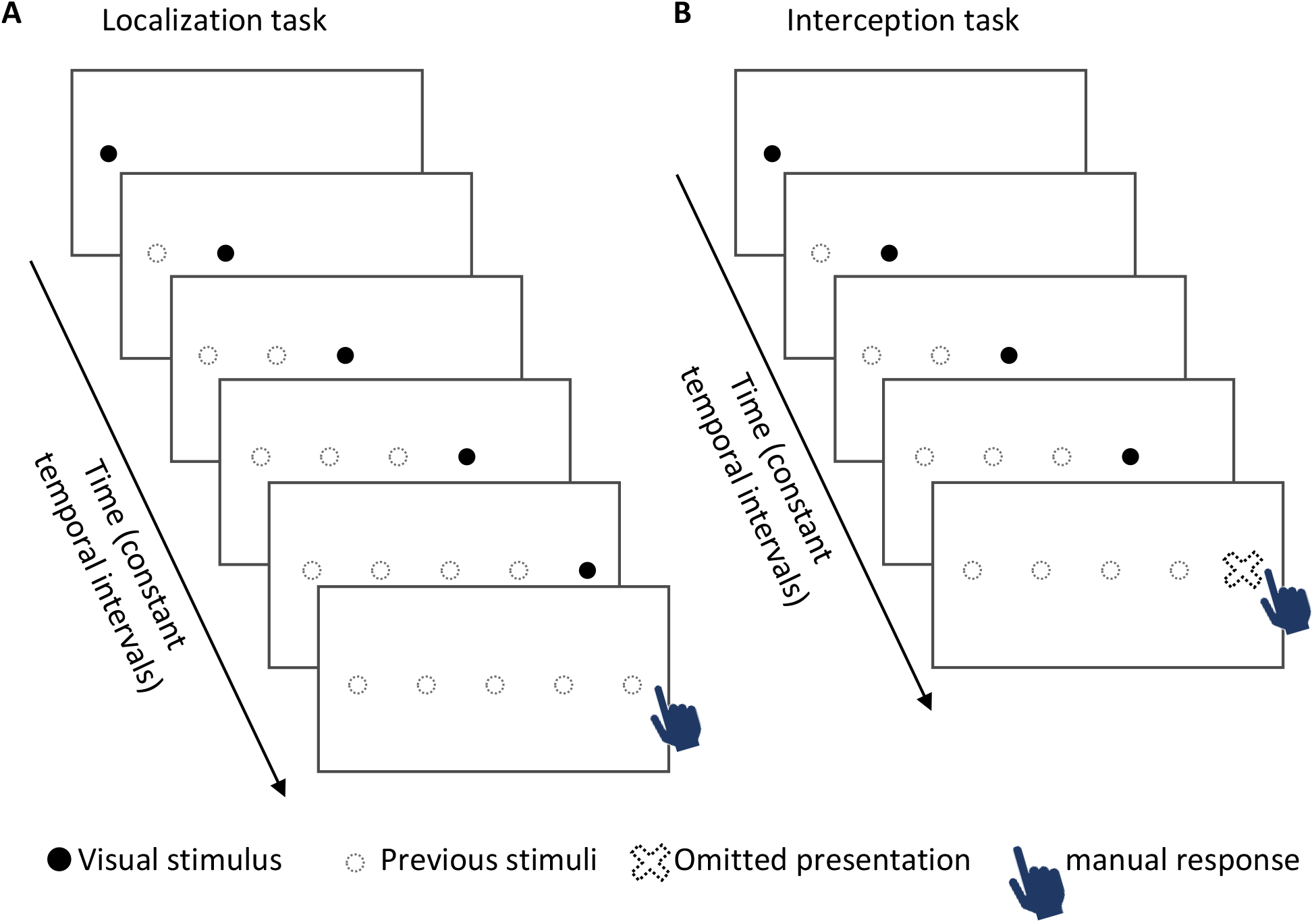
Experimental Procedure of localization (A) and interception (B) paradigms. A black dot was intermittently presented on either 5 (A) or 4 (B) locations with constant spatial and temporal interstimulus intervals. Participants had to either indicate the remembered final location after stimulus presentation (A) or predict the time and location of the (omitted) fifth presentation (B). ‘Movement’ direction (left-right vs. right-left) was randomized.

Each trial was programmed in a way that the fifth location was randomly chosen from a range of -100 px to 100 px (vertically as well as horizontally) from the center of the screen. This procedure induced uncertainty about the correct final location while ensuring that the target ends within reaching distance of the participants.

Overall, participants worked through 8 practice and 144 experimental trials per block (3 temporal ISI x 3 spatial ISI x 2 directions x 8 repetitions), resulting in 304 trials in total. Practice trials were identical to the upcoming experimental block but only consisted of the two most extreme temporal and spatial interstimulus intervals.

### Statistical Analysis

To compare biases in both tasks, we fitted two linear mixed models with trials nested in participants to the horizontal localization and interception errors (overshooting), respectively. Each model included fixed and random effects of temporal and spatial interstimulus intervals, as well as random intercepts to predict the horizontal error. From each model, the intercepts and slopes per participant were extracted. Both interstimulus intervals were scaled (mean = 0, sd = 1), so that the intercept reflects the average overshooting bias, whilst the slope indicates how strongly temporal manipulations impact this bias. To evaluate whether the amount of overshooting generalizes across tasks, we correlated the two intercepts. Additionally, we correlated the individual slopes between the two tasks to estimate whether manipulations of temporal features similarly affect the bias in spatial representations (localization task) as well as spatiotemporal prediction (interception task). Given that the correlations are limited by reliabilities, we additionally calculated the mean split-half reliabilities of each scale (out of 20 random samples) and corrected the correlations by dividing them by the square root of the multiplied reliabilities (Murphy & Davidshofer, 1988).

## Results

Participants on average missed the correct location by 0.16 cm in the localization task and by 1.01 cm in the interception task. When intercepting they reached the screen approximately 600 ms later than the omitted target. In some trials during the localization task, participants released the space bar too early, that is, even before the fifth stimulus presentation started (22 trials in total). These trials were excluded for further analysis. Outliers for each participant in each task and spatial or temporal condition (3 sd below or above the mean) and trials in which participants responded more than 3s too late were additionally excluded. This resulted in 0.9% excluded data in the localization task and 0.3% in the interception task.

### Effect of temporal intervals on localization and interception

Two linear mixed models predicting the amount of overshooting from the temporal intervals revealed a positive intercept and a significant negative effect of temporal intervals on the horizontal touch location. **Figure 2A** shows the horizontal error of four illustrative participants (ids: 5, 20, 59, 63) for the localization (dark red) and the interception task (bright red) across the three temporal intervals. Overall, participants overshot the correct location in the localization task, intercept = 0.16, *t*(66) = 6.93, *p* < .001, as well as the interception task, intercept = 1.01, *t*(66) = 10.15, *p* < .001. This overshooting decreased with increased temporal intervals (i.e., slower speed) when participants indicated the remembered position (localization), β = -0.04, t(66) = -3.49, *p* < .001, and also when they predicted the future location (interception), β = -0.31, *t*(65.9) = -10.20, *p* < .001. Spatial intervals only affected localization accuracy, β = -0.03, *t*(66.1) = -3.30, *p* = .001, but not interception accuracy (*p* = .651). With longer spatial intervals, participants overshot the final location less in the localization task.

**Figure 2.**
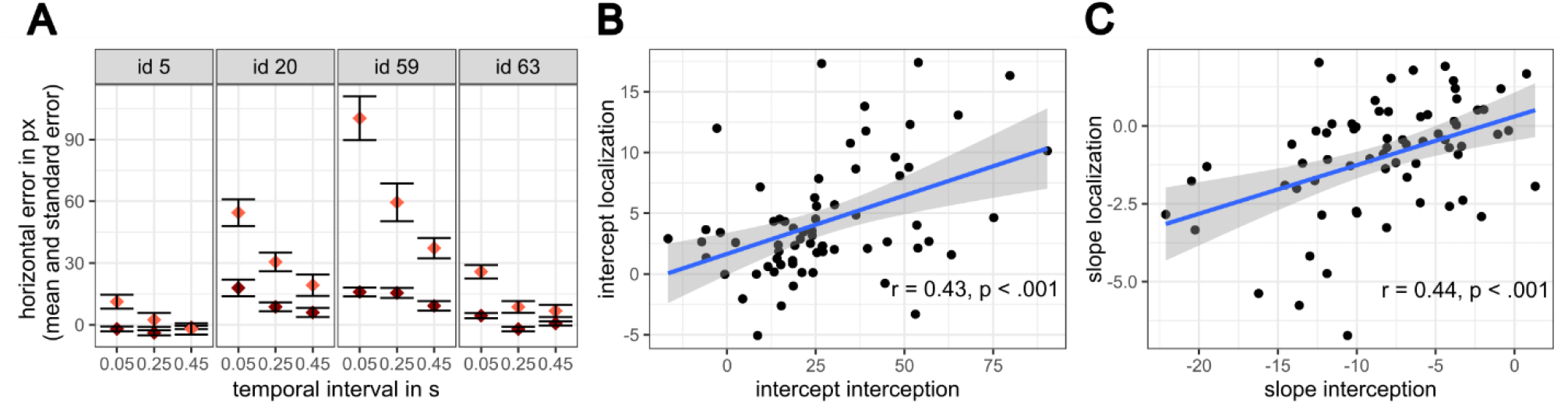
Localization and interception errors. **A)** Four illustrative participants showing the overall overshooting effect in localization (dark red) and interception (bright red) decreasing with increasing temporal intervals. Positive values indicate errors in motion direction. **B)** Correlation of the overall overshooting effects drawn from the intercepts of the linear mixed models. Participants who overshoot in localization also overshoot in interception. **C)** Correlation of the temporal manipulation effects drawn from the slopes of linear mixed models. If a participant is largely affected by the temporal manipulation in one task, this participant is also largely affected in the other task.

### Association between localization and interception

To test for an association between the overshooting in both tasks, we correlated the intercepts of the model fits of each participant across tasks. This revealed a positive correlation, indicating that participants who showed larger overshooting for the remembered fifth location (localization) similarly showed larger overshooting when predicting this location (interception), *r*(65) = 0.43, reliability-corrected *r* = .46, *p* < .001 (**Fig. 2B**). Split-half reliabilities were very high for both intercepts (localization: mean r = .91, sd = .02, interception: mean r = .96, sd = .01). Additionally, we tested whether these spatial biases change proportionally with increasing temporal intervals across participants. Indeed, participants with larger slopes in the localization task also expressed larger slopes in the interception task, *r*(65) = 0.44, reliability-corrected *r* = .65 *p* < .001 (**Fig. 2C**). Split-half reliabilities were moderate to high for both slopes (localization: mean r = .71, sd = .05, interception: mean r = .64, sd = .08). Both effects indicate that performance in localization and interception is interrelated.

## Discussion

We tested whether spatial biases of moving objects observed in two areas of research, namely localization (measured by the representational momentum) and interception (measured by the tau effect), are related to each other. Participants either indicated the remembered final location or predicted a future location of an intermittently presented moving (‘jumping’) dot by tapping on a touchscreen. We showed that i) the amount of overshooting in localization was closely related to the amount of overshooting in interception across participants, and that ii) participants were similarly biased by temporal manipulations in both tasks, with shorter ‘jump times’ leading to more overshooting. Both results indicate that errors in localization and interception are related to each other and might suggest that these biases are driven by the same underlying mechanism.

### Motion perception, temporal prediction and spatiotemporal action

A similar approach of leveraging individual differences has already been applied to show that the shift in localization predicts time-to-contact estimates (Gray & Thornton, 2001).

Extending these findings, we now additionally show that these shifts also predict how much participants overshoot in interception. All these results imply a general bias in motion processing which does not only emerge in perceptual performance but also in motion predictions and motor actions (see also, Schroeger, Raab, et al., 2022). On a broader scale these results fit in the line of research showing a transfer of perceptual biases to action tasks (e.g., Franz et al., 2000; Rossetti et al., 2017).

We did not only rely on individual differences in the overall bias, but also applied experimental manipulations in both tasks. Even though the actual location was independent of the temporal intervals, our temporal manipulation still affected both, overshooting in localization and interception. Again these results indicate that spatial judgements are not only related to temporal (time-to-contact estimates, Gray & Thornton, 2001) but also spatiotemporal predictions (interception). The question arises as to why temporal information affects the spatial responses. Several theories have suggested that spatial and temporal representations are closely interrelated and therefore impact each other (e.g., a theory of magnitude, Walsh, 2003; Cai & Wang, 2022; conceptual metaphor theory, Lakoff & Johnson, 1980). Specifically in the context of motion, space and time are typically highly correlated, mediated via the movement speed. Accordingly, participants might implicitly expect that temporal changes are associated with changes in speed and therefore impact the final location, even if they were disentangled in the current task.

Overall, our results show that localization and interception might largely rely on similar processes and therefore be subject to the same biases.

### Potential underlying mechanisms

Several theories attempt to explain these effects. One prominent idea posits that a (slow) speed prior causes errors in motion perception and prediction (Goldreich, 2007; Goldreich & Tong, 2013; Merz et al., 2022; Weiss et al., 2002). This suggests we even infer motion from sequentially presented stationary stimuli (Goldreich, 2007), a concept distinct from fluent apparent motion (Wertheimer, 1912). This speed prior, used to reduce uncertainty in inherently uncertain motion perception, leads to biased percepts. If a shared underlying mechanism, such as a speed prior, drives these perceptual biases, we would expect correlated effects. Indeed, this study empirically demonstrates that individuals showing a larger representational momentum effect also exhibit a larger tau-interception effect. Furthermore, manipulating temporal intervals similarly affects both biases, supporting the notion of similar mechanisms, potentially speed expectations, to underlie motion perception. However, Kirsch (2023) recently proposed a Bayesian cue integration model, without relying on a speed prior, where spatial and temporal magnitude information interact to explain both the tau and kappa effects.

Similar to previous research, our findings also indicate individual differences in these biases (Gray & Thornton, 2001), presenting an opportunity to relate them and explore potential underlying factors (Goettker & Gegenfurtner, 2024). These variations might stem from different speed priors developed through experiences and adaptation to the experimental context (Merz et al., 2023), potentially explaining various spatial and temporal biases in motion perception, including effects like the Fröhlich effect (Fröhlich, 1923). Consequently, a broader investigation of multiple motion processing effects could lead to a more comprehensive understanding and potentially reveal the role of speed priors for motion perception. While the original tau paradigm involved different task demands compared to our motion prediction and interception tasks, examining correlations across these could help differentiate task-related and (potentially) speed-prior-related shared variance.

## Conclusion

We found an association between two spatial biases reported in the literature, one in a localization (representational momentum) and one in an interception (tau) task. This was not only revealed by the overall amount of overshooting but also by the way in which this overshooting was affected by temporal manipulations. Our results indicate that motion perception/localization and motion prediction/interception share similar biases and subsequently likely underlying processes.

## Acknowledgements

We thank Marie Mengel for her help with data collection.

## Data availability statement

Data and Materials will be made public on osf upon acceptance for publication.

## Notes

A.S. was supported by the Research Cluster “The Adaptive Mind”, funded by the Excellence Program of the Hessian Ministry of Higher Education, Science, Research and the Arts. S.M. was supported by a grant of the German research foundation (DFG) under grant number ME 5568/1-2.

### Competing Interest Statement

The authors have declared no competing interest.

